# Hybrid EEG-EMG system to detect steering actions in car driving settings

**DOI:** 10.1101/2021.09.16.460615

**Authors:** Giovanni Vecchiato, Maria Del Vecchio, Jonas Ambeck-Madsen, Luca Ascari, Pietro Avanzini

**Affiliations:** Institute of Neuroscience, National Research Council of Italy, Parma; Institute of Neuroscience, National Research Council, Italy, Parma; Toyota Motor Europe; Camlin Italy s.r.l.

**Keywords:** hybrid system, driving, steering, EEG, EMG, time-frequency analysis

## Abstract

Understanding mental processes in complex human behaviour is a key issue in the context of driving, representing a milestone for developing user-centred assistive driving devices. Here we propose a hybrid method based on electroencephalographic (EEG) and electromyographic (EMG) signatures to distinguish left from right steering in driving scenarios. Twenty-four participants took part in the experiment consisting of recordings 128-channel EEG as well as EMG activity from deltoids and forearm extensors in non-ecological and ecological steering tasks. Specifically, we identified the EEG mu rhythm modulation correlates with motor preparation of self-paced steering actions in the non-ecological task, while the concurrent EMG activity of the left (right) deltoids correlates with right (left) steering. Consequently, we exploited the mu rhythm de-synchronization resulting from the non-ecological task to detect the steering side by means of a cross-correlation analysis with the ecological EMG signals. Results returned significant cross-correlation values showing the coupling between the non-ecological EEG feature and the muscular activity collected in ecological driving conditions. Moreover, such cross-correlation patterns discriminate left from right steering with an earlier dynamic with respect to the single EMG signal. This hybrid system overcomes the limitation of the EEG signals collected in ecological settings such as low reliability, accuracy and adaptability, thus adding to the EMG the characteristic predictive power of the cerebral data. These results are a proof of concept of how it is possible to complement different physiological signals to control the level of assistance needed by the driver.

## 1. Introduction

Recent advances in sensing and control techniques have made it possible to design cars with a large set of features which can take control of specific aspects of the driving task (e.g cruise control, autonomous driving) or provide information to the driver to support specific manoeuvres (e.g. lane departure). Because driving is a complex behaviour involving interrelated motor and cognitive elements such as attention, visuospatial interpretation, visuomotor integration and decision making (Calhoun et al. 2002; Calhoun and Pearlson 2012), to achieve ever higher levels of driver support it is important to investigate and characterize the related brain processes underlying the driver’s actions.

In the years, several attempts were made to characterize the neural correlates of driving in scenarios with changing level of ecology and different neuroscientific technique investigating haemodynamic (Walter et al. 2001; Spiers and Maguire 2007; Mader et al. 2009; Calhoun and Pearlson 2012; Schweizer et al. 2013), magneto- (Fort et al. 2010; Sakihara et al. 2014) and electrophysiological (Schier 2000; Haufe et al. 2011, 2014; Gheorghe et al. 2013; Khaliliardali et al. 2015; Kim et al. 2015; Zhang et al. 2015; Brooks and Kerick 2015; Brooks et al. 2016; Garcia et al. 2017; Vecchiato et al. 2018, 2020) activity. In particular, these last works show the possibility to use electroencephalography (EEG) to decode the driver’s cognitive processes in simulated and real car scenarios with the ultimate goal to predict the upcoming action. Despite the success in the classification of salient driving events such as braking (Haufe et al. 2011, 2014; Kim et al. 2015) and steering (Gheorghe et al. 2013) actions, the level of accuracy is still moderate and, most importantly, the detected neurophysiological features are elicited just around few milliseconds before the upcoming driving event making it difficult the future implementation of electronic assisting devices. Moreover, acting in realistic scenarios means taking into account the variability of a dynamic environment leading to the complexity of human behaviour which arise from a variety of mental processes acting in parallel producing high variability in the EEG. This motivates the need to identify other robust physiological features tackling increased noise due to environmental characteristics and to the interaction among neural processes (Lohani et al. 2019).

In this sense, surface electromyography (EMG) provides a non-intrusive way of measuring muscle activation and is an appropriate technique when assessing active steering systems (De Luca 1997; Ahlström et al. 2019). It opened new perspectives in the field of ergonomics and provides new tools for the analysis of the neuromuscular system in working environments. It is experiencing a growing interest in medical and research applications thanks to the recent availability of novel low-end commercial products increasing the wearability with respect to EEG sensors allowing to perform longer recording sessions in a more comfortable way (Gazzoni et al. 2016; Milosevic et al. 2017). Previously, EMG recordings have been used to assess the function of the muscles of the upper limb in car driving (Jonsson and Jonsson 1975; Liu et al. 2012; Gao et al. 2014). The main findings are that the prime movers are primarily a consequence of steering direction while the stabilizing or fixating muscles are primarily constant, returning that the key muscles correlated to steering are the triceps brachii, the deltoids, pectoralis major and infraspinatus (Pick and Cole 2006; Liu et al. 2012; Gao et al. 2014). In particular, it is well known that the whole deltoid muscle acts in abduction of the arm and there is a synergy between the anterior portion of the muscle and the contralateral posterior portion when moving the steering wheel. More specifically, the anterior portion serves to rotate the steering wheel contralaterally and the posterior portion to rotate it ipsilaterally (Jonsson and Jonsson 1975). However, despite high values of signal-to-noise-ratio and ergonomic advantages, the EMG signal is characterized by poor anticipation power in predicting the upcoming action because it results the last element of the neuromuscular chain.

It has been demonstrated that voluntary movements generate changes in the alpha and beta ranges of the EEG spectrum (Pfurtscheller and Lopes da Silva 1999), and that such rhythms are in turn related to modulations of the EMG activity (Hari and Salenius 1999). Recent works have focused on the synchronization between rhythmical activity in the motor cortex and muscular activity employing cortico-muscular coherence, which is usually observed during periods of muscular contraction, and has been reported in a number of studies involving both EEG and MEG (Conway et al. 1995; Boonstra et al. 2009; Cheyne 2013; Rizzo et al. 2020). Current approaches to cortico-muscular coordination focus on associations and synchronous activation between individual brain rhythms at specific cortical areas (e.g., motor cortex, hyppocampus), and peripheral muscle activity during specific movement tasks or exercises in ecological conditions (walking, running, etc.) (van Wijk et al. 2012; Rendeiro and Rhodes 2018). In such scenarios the neuromuscular signals are noisier than the ones collected in standard laboratory conditions due to uncontrolled environmental settings, and there is also the issue to address the simultaneous presence of concurrent cerebral processes. Finally, on the user side there is the need to create comfortable experimental settings to not pollute the neuromuscular signals with undesired components due to the experienced fatigue of wearing biomedical sensors on the head and on several part of the body. To overcome issues with both EEG-and EMG-based control methods, a combination of both systems, building on the advantages of each signal and diminish the limitations of each might be a promising strategy (Lalitharatne et al. 2013). On this research line, it would be useful to identify cerebral features of interest in standard laboratory settings and then looking for neuromuscular invariants in more ecological settings characterized by a lower recording complexity and highest comfort. In this way, off-line analysis performed on data collected in standard settings would be informative of the on-line pipeline to implement to extract neuromuscular features of interest.

Following this reasoning, here we propose a hybrid method to distinguish left from right steering in driving simulator based on EEG and EMG signals collected during steering actions. In particular, we aim to identify EEG features of movement preparation which would anticipate the EMG activity leading to the action onset. The combination of the two signals would increase the predictive power of the steering action. We investigated brain and muscular activity underlying steering behaviour during a non-ecological steering task. Such paradigm allowed us to identify electrophysiological correlates of steering preparation without confounding effect. Then, we correlated such features with the EMG activity collected during a session of driving simulation to extend the non-ecological cerebral signatures to a more ecological steering task. Therefore, we aimed to characterize the EEG-EMG coupling associated with the natural and self-initiated execution of steering actions while driving. To this end, we took advantage of the Independent Component Analysis (ICA) which separates mixed EEG signals into maximally independent activities, each characterized by a precise scalp topography and a corresponding generator pattern, typically modelled as patches of cortical pyramidal cells (Delorme et al. 2012). This strategy also allowed us to avoid field spread caused by the large distance between sensors and neural sources and by the spatial blurring effect of the skull on the scalp’s potential distribution of EEG signals (Schoffelen and Gross 2009). Thus, we expect to identify reliable independent components, whose topography indicates the involvement of motor circuits, whose reactivity precedes the steering action and is correlated with the muscular activity of the deltoids. Should the EEG-EMG coupling be modulated across the two types of action performed by participants (i.e., left and right steering), this would provide evidence that such hybrid method can be used to discriminate and possibly predict motor intentions associated with natural driving behaviour.

## 2. Material and methods

### 2.1. Participants

Twenty-four participants (6 females, M: 22.8 ± SD: 2.0 years of age) took part in this experiment. They verbally declared that they were right-handed and had normal or corrected to normal vision, no history of neurologic or psychiatric disorders and no daily medications. All of them have driving license obtained from the Italian state authority for motor vehicles. None of them had driving experience on professional racetracks. They gave written informed consent to participate in the study consisting into two consecutive sessions comprising a non-ecological steering task and a following ecological steering task. Approval for the study was obtained by the local Ethical Committee (comitato etico Unico per la provincia di Parma).

### 2.2. Experimental scenarios

#### Non-ecological steering task

Participants seated in front of a computer screen at a distance of 1 m and were instructed to turn at a quiet pace the steering wheel (Logitech G25) on the right or on the left according to a traffic sign randomly presented on the display after 1 s from an attentional cue (i.e., a white cross). Participants who performed the steering within 2 s from the traffic sign received an error message and repeated the trial. According to this rule, 88 correct trials have been collected for each participant, equally distributed between left and right steering actions. Synchronization between the visual stimulation, participant’s behavior and EEG data was realized by using timestamps sent by a photodiode placed in front of the screen capturing the task events through white labels onto a black background. Figure 1 shows the experimental setup (panel A), and the time-course of the task (panel B).

**Fig.1.**
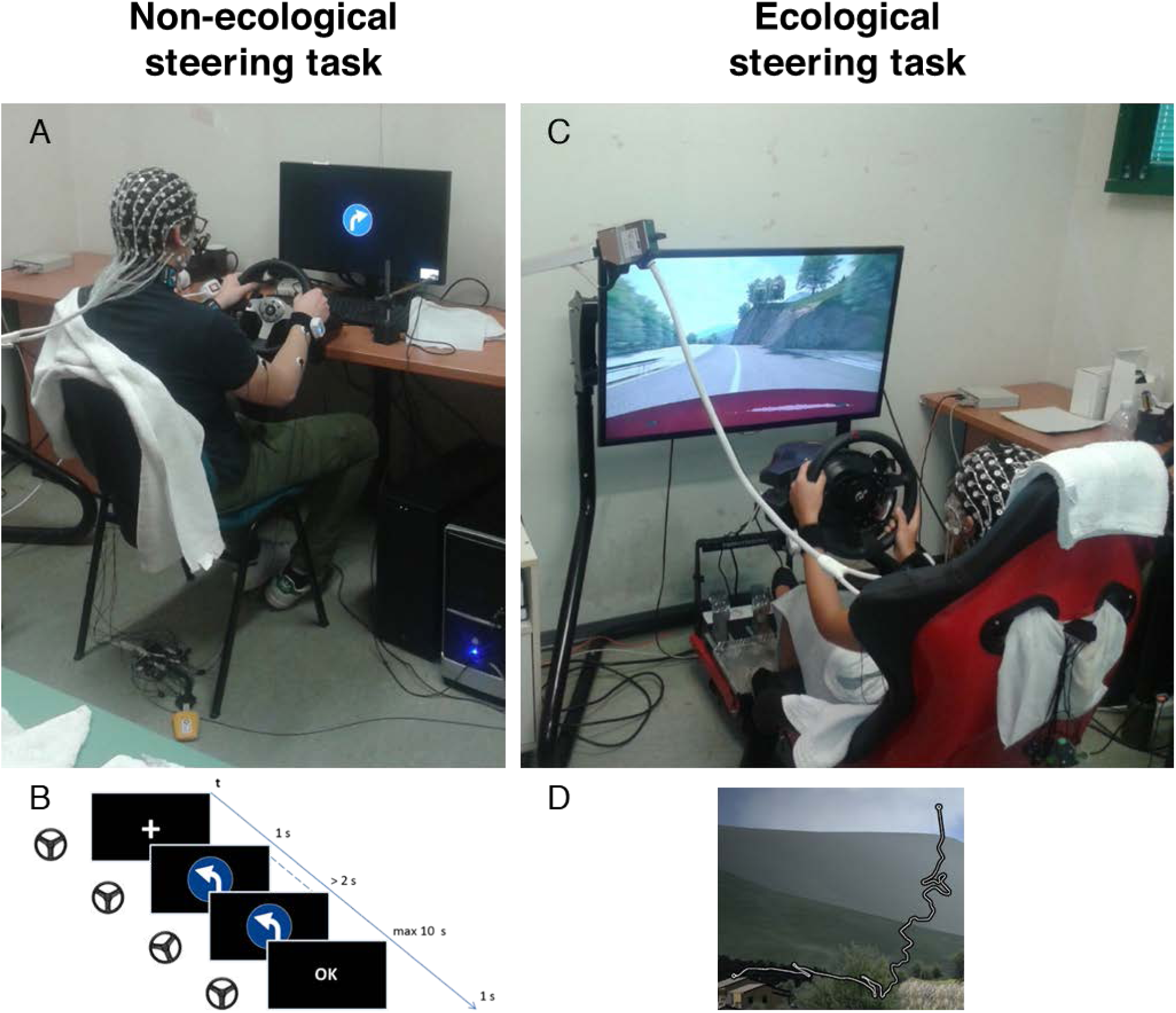
Pictures of the experimental setups related to the non-ecological (panel A) and ecological (panel C) steering task. Panel B shows the timeline of the events during the non-ecological task. Panel D presents the road profile of the track used in the ecological steering task

#### Ecological steering task

Participants seated in a driving simulator composed of the RSeat RS1 Assetto Corsa Special Edition (seat) and Thrustmaster T500 RS (steering wheel and pedals). A video game (Assetto Corsa - Kunos Simulazioni) was displayed on a Samsung 40’’ 5300 class LED TV positioned 1 m from the participant’s seat (vertical field view = 27.4°, horizontal field view = 46.8°). The car used for the experiment was an Alfa Romeo Mito with an automatic transmission. First, participants were asked to get familiar with the environment and the setup by driving for a lap on the Monte Erice track (downloadable at http://assettocorsa.club/mods/tracks/monte-erice.html). Afterwards, they performed a single lap on the Coste Loop track (part of the Assetto Corsa software), maintaining the right lane with no particular constraints. They were asked to drive as if it was their own car in a natural manner. This circuit simulates a part of the Garda Lake coastal road. No other vehicle was present on both tracks. During the experiment, we collected data related to the steering wheel angle. The synchronization among all the recording devices was implemented with the Lab Streaming Layer (LSL) as described in our previous work (Vecchiato et al. 2018). Figure 1 shows the experimental setup (panel C), and a representation of the Coste Loop track (panel D).

### 2.3. Behavioural data collection and analysis

The steering wheel signals was collected during both the non-ecological and ecological tasks and were segmented in trials [-2000, 2000] ms around the steering onset. In order to identify the steering onset in the non-ecological task, we consider the first part of the trial returning wheel angles above the threshold of ±2°. Of this segment, the time bin corresponding to the point of maximum distance computed from the steering curve and the segment joining the first and the last sample of the trial was identified as the steering onset (see Supplementary Material for a schematic representation of the procedure). When an uncertainty in the movement onset arose, we discarded the trial (around 3.5%). The result of this procedure was also used to epoch the EEG and EMG signals collected during the non-ecological task into [-2500, 2500] ms trials around the steering onset.

Instead, in the ecological task the steering onset was determined through a procedure of quantization of the steering wheel angle signal. The quantization was realized partitioning the wheel signal with a step of 0.125 between its minimum and maximum. This step was set to assure that the error between the areas calculated from the original and the quantized signals was below 5%, and the correlation between the two was greater than 99%. The quantized signal was binarized setting at 1 all values greater than 0 (in absolute value) and then clustered. The steering onset was determined as the time bin corresponding to the transition between zeros and ones. With such procedure, we identified 94.58 ± 31.12 right steering and 79.08 ± 23.85 left steering trials.

Moreover, the steering wheel signal collected during the ecological task was also segmented during the non-steering intervals in [-2000, 2000] ms overlapping trials of 1 second, thus identifying 163.71 ± 103.55 trials.

### 2.4. EEG and EMG data recording and pre-processing

Continuous EEG was recorded in the non-ecological task using the 128-channel Geodesic EEG System (Electrical Geodesics, Inc., Eugene, OR, USA) and the HydroCel Geodesic Sensor Net. Consistent positioning was achieved by aligning the Sensor Net with skull landmarks (nasion, vertex and pre-auricular points). Using high-input impedance amplifiers (Net Amps300), low-noise EEG data was obtained with sensor-skin impedances maintained below 50 kΩ. The signal was digitised at a sampling rate of 500 Hz (0.01 Hz high-pass filter) and recorded with a vertex reference, the impedance of which was kept below 10 kΩ. Impedances were checked and adapted newly at the beginning of the ecological steering task. EEG data were exported in raw format using NetStation software (Electrical Geodesics, Inc., Eugene, OR, USA) and then imported into MATLAB to perform the following analysis with EEGLAB v14.1.2 (Delorme and Makeig 2004). Pre-processing comprised line noise removal, bad channels interpolation and common average reference. Since the steering action was a self-paced movement many trials lasted more than 10 seconds. Hence, EEG data were segmented into epochs [-1500, 11500] ms around the presentation of the steering traffic sign to take into account even the longest trials. Artefacts were rejected by applying a semi-automatic procedure to detect abnormal trends and spectra. On average, we discarded 11.4 ± 7 trials. Clean EEG datasets were comprised of 37.6 ± 3.7 left and 37.7 ± 3.3 right steering trials. Then, we performed an independent component analysis (ICA) to identify and separate neurophysiological brain activities from other noise sources. On average, we identified 5.9 (± 2.1) independent components (ICs) per subject, for a total sum of 147 EEG ICs. Cluster analysis was performed to group components according to their scalp topographies via the K-means algorithm (Lloyd 1982). The number of clusters was chosen by a bootstrap-based method which lead to the computation of a stability index (Ben-Hur et al. 2002; Salvador and Chan 2004) and the identification of 6 clusters. Details related to the pre-processing pipeline, independent component analysis and clustering can be found in our previous work (Vecchiato et al. 2018).

Continuous EMG data in both non-ecological and ecological steering tasks were acquired using the Neuroelectrics Enobio. EMG signals were sampled at 500Hz from the left and right deltoids, left and right forearm extensor digitorum as among the main muscles involved in steering actions (Pick and Cole 2006; Lohani et al. 2019) and later imported and processed in MATLAB environment (R2018b, The Mathworks, Natick, MA). EMG data of non-ecological task were segmented into epochs [-1500, 11500] ms around the presentation of the steering traffic sign (as done for the EEG data), while EMG data of the ecological task were segmented into two datasets: i) epochs [-2500, 2500] ms around the onset of the steering action, as well as ii) 1 s overlapping epochs of [-2500, 2500] during non-steering intervals. Line noise of the first 5 harmonics of 50 Hz was suppressed using a spectrum estimation technique (Mewett et al. 2001).

### 2.5. EMG and EEG data analysis

For each EMG signal and EEG IC, we computed the event-related spectral perturbation (ERSP) as time-frequency decomposition using Morlet wavelets (Makeig et al. 2004). For the EMG data we used frequencies that increased from 2 to 200 Hz in 198 linearly spaced steps, with the number of wavelet cycles increasing from 3 to 60 in linear steps. For the EEG data we used frequencies that increased from 2 to 40 Hz in 38 linearly spaced steps, with the number of wavelet cycles increasing from 3 to 12 in linear steps. Then, dB conversion was performed with a single-trial baseline normalization using the [-500, -100] ms time window before the steering sign presentation for the non-ecological steering task, and the interval of [-2000, 2000] ms for the ecological paradigm.

To identify the possible time-lag between the EEG signals recorded in the non-ecological task and the EMG signal recorded in the ecological task, we computed a cross-correlation analysis between the corresponding time-frequency panels. In particular, for each subject and steering condition in the non-ecological task, we extracted an EEG mask segmenting the time-frequency panel around [8, 20] Hz and [-1500, -1000] ms before the steering onset (e.g., left non-ecological steering). Then, for each subject we performed the bi-dimensional cross-correlation trial-by-trial between such EEG mask and the whole EMG time-frequency panel (e.g, left ecological steering). The specificity of the resulting cross-correlation patterns with respect to the steering action was addressed by computing the same analysis using the EMG signals related to non-steering intervals. In particular, these data were randomly assigned to left and right pseudo-steering conditions with a half split technique: for each of 300 iterations non-steering trials were divided in two sets, only one of this subset was used to create the left and right pseudo-steering conditions. Then, for each iteration we computed the cross-correlation values with the EEG mask resulting from the analysis of the non-ecological dataset. Finally, we also tested the specificity of the cross-correlation patterns with respect to the directionality of the steering by adopting a shuffling procedure: for each of 300 iterations we created the pseudo-left and pseudo-right conditions by randomly assigning trials from the original left and right steering datasets and then computing the cross-correlation values.

To discriminate the EEG and EMG activity between steering sides (i.e., left vs right steering) and EMG recording sites (i.e., left vs right deltoid) in both non-ecological and ecological tasks, as well as the cross-correlation signals computed in the ecological task, the corresponding time-frequency panels were compared using dependent sample t-statistics and non-parametric permutation testing, corrected for multiple comparisons by weighted cluster mass correction with randomisation of 1000 and a statistical threshold of 0.05 (Hayasaka and Nichols 2004; Maris and Oostenveld 2007). The significance of the t-statistics computed to test the specificity for steering and directionality was assessed by comparing the observed statistics to the statistical properties of the null-hypothesis distribution. Hence, the observed test statistic values were converted into Z scores and then the corresponding p-values were computed and reported (Cohen 2014).

## 3. Results

### 3.1. Non-ecological steering task

For each cluster of EEG ICs, we performed the non-parametric permutation test to compare the ERSP between the two steering conditions (i.e., left vs right) in the time window of [-2000, 2000] ms segmented with respect to onset of the event. The analysis revealed a statistically significant difference onset only for cluster 3, which highlights a desynchronization of the mu rhythm around 1500 ms before the left steering. This cluster is populated by 24 ICs belonging to 16 subjects. Figure 2 shows the scalp topography of such significant cluster along with the modulation of the related time-frequency activity during left (panel A) and right (panel B) non-ecological steering with the corresponding t-statistics (panel C). Centroid maps with the corresponding time-frequency panels of the non-significant EEG IC clusters are illustrated in the Supplementary Material.

**Fig.2.**
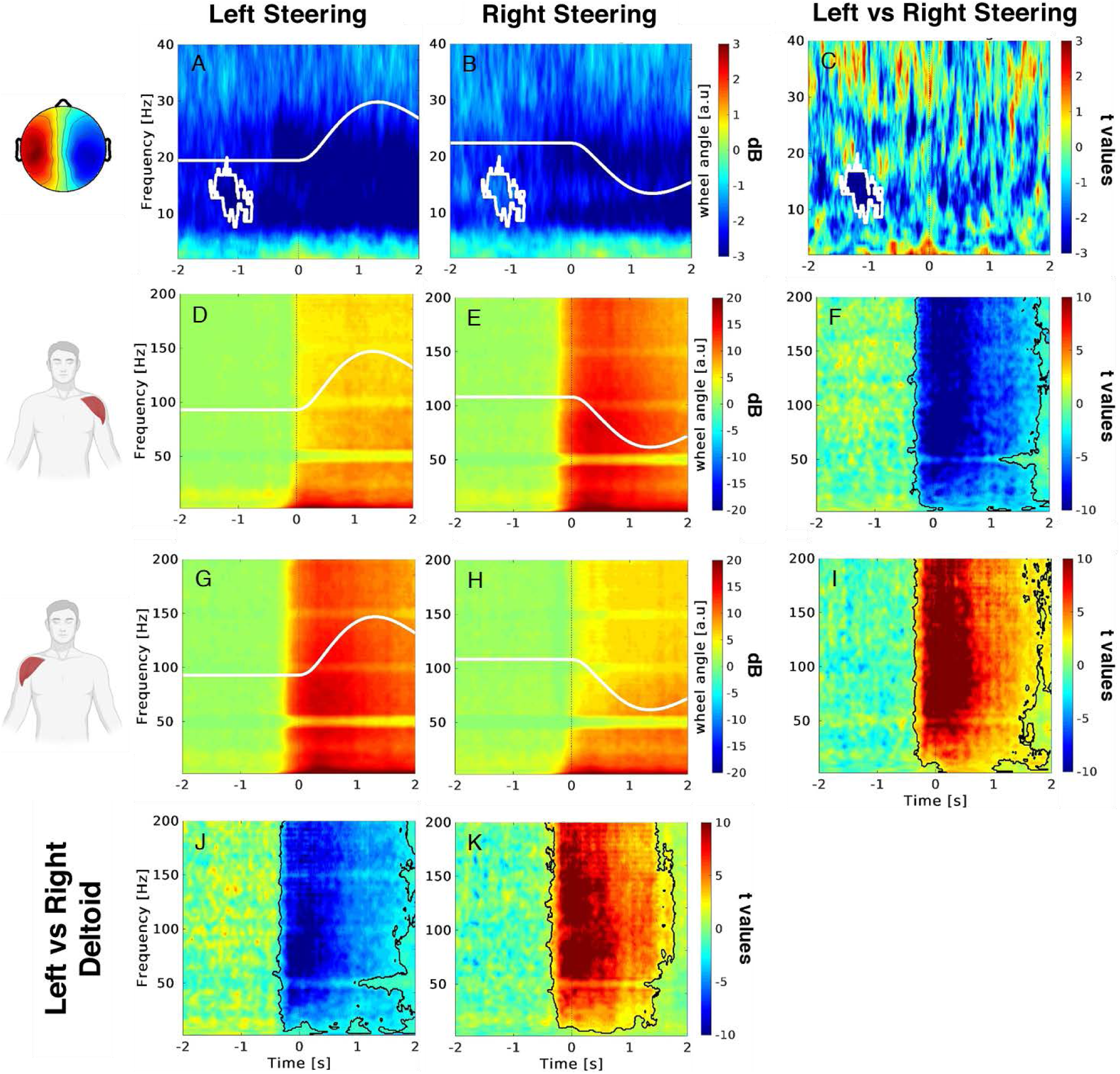
ERSP for the EEG and EMG signals collected during the non-ecological steering task. Upper row illustrates the EEG ERSP for left (A) and right (B) steering as well as the statistical comparison of the two conditions (C). The topography in the left part of the picture shows the average scalp map related to the cluster 3 centroid. Second row (from the top) illustrates the EMG ERSP for the left deltoid during left (D) and right (E) steering as well as the statistical comparison of the two conditions (F). Third row illustrates the EMG ERSP for the right deltoid during left (G) and right (H) steering as well as the statistical comparison of the two conditions (I). Lower row illustrates the statistical comparisons of the EMG ERSP between left and right deltoid during left (J) and right (K) steering. Colorbars of the upper row indicate in blue (red) the desynchronization (synchronization) of the EEG activity (A, B) with respect to the baseline, as well as the statistical differences corresponding to the increase (decrease) of such activity during the left (right) steering (C). Colorbars of the second and third rows indicate in red (blue) the synchronization (relaxation) of the EMG activity (D, E, G, H) with respect to the baseline, as well as the increase (decrease) of such activity during the left (right) steering for the right (left) deltoid, as well as for the left (right) deltoid during right (left) steering (F, I, J, K). White lines depict the left and right steering wheel angle profiles (A, B, D, E, G, H). White mask (A, B, C) delimits the statistically significant portion of the EEG ERSP panel. Black mask (F, I, J, K) delimits the statistically significant portion of the EMG ERSP of the corresponding compared panels (non-parametric t-test, cluster corrected)

Figure 2 also shows the ERSP of the EMG activity for the left (panels D, E) and right deltoids (panels G, H) during the left (panels D, G) and right (panels E, H) steering. Specifically, when comparing these signals between left and right steering, we observed the significant broadband increase of the muscular activation of the left deltoid during the right steering (panel F), as well as the symmetrical broadband increase of activity of the right deltoid during the left steering (panel I). Analogously, when comparing these signals between left and right deltoids, we observed the significant broadband increase of the muscular activation of the right deltoid during the left steering (panel J), as well as the symmetrical broadband increase of activity of the left deltoid during the right steering (panel K). The statistical comparisons of the muscular activation of the two forearm extensors did not return significant differences in the EMG broadband as the two deltoids (see Supplementary Material, Figure S3), thus they are not considered in the following analysis.

Variations of the wheel angle for the left and right steering were represented with the white signals within each corresponding panel of Figure 2.

### 3.2. Ecological steering task

Figure 3 shows the results related to the analysis of the EMG signals during the ecological task performed at the driving simulator. In particular, the upper box presents the ERSP computed for the EMG signals of the left and right deltoids during left and right steering actions. The different panels of this figure are arranged as the Figure 2 to show that the asymmetrical activation of the two deltoids during steering observed in the non-ecological task is replicated in the ecological condition, too. Specifically, we highlight the significant broadband increases of the muscular activation of the left deltoid during the right steering (panels A-C), as well as the symmetrical broadband increase of activity of the right deltoid during the left steering (panels D-F). Analogously, when comparing these signals between left and right deltoids, we observed the significant broadband increase of the muscular activation of the right deltoid during the left steering (panel G), as well as the symmetrical broadband increase of activity of the left deltoid during the right steering (panel H).

**Fig.3.**
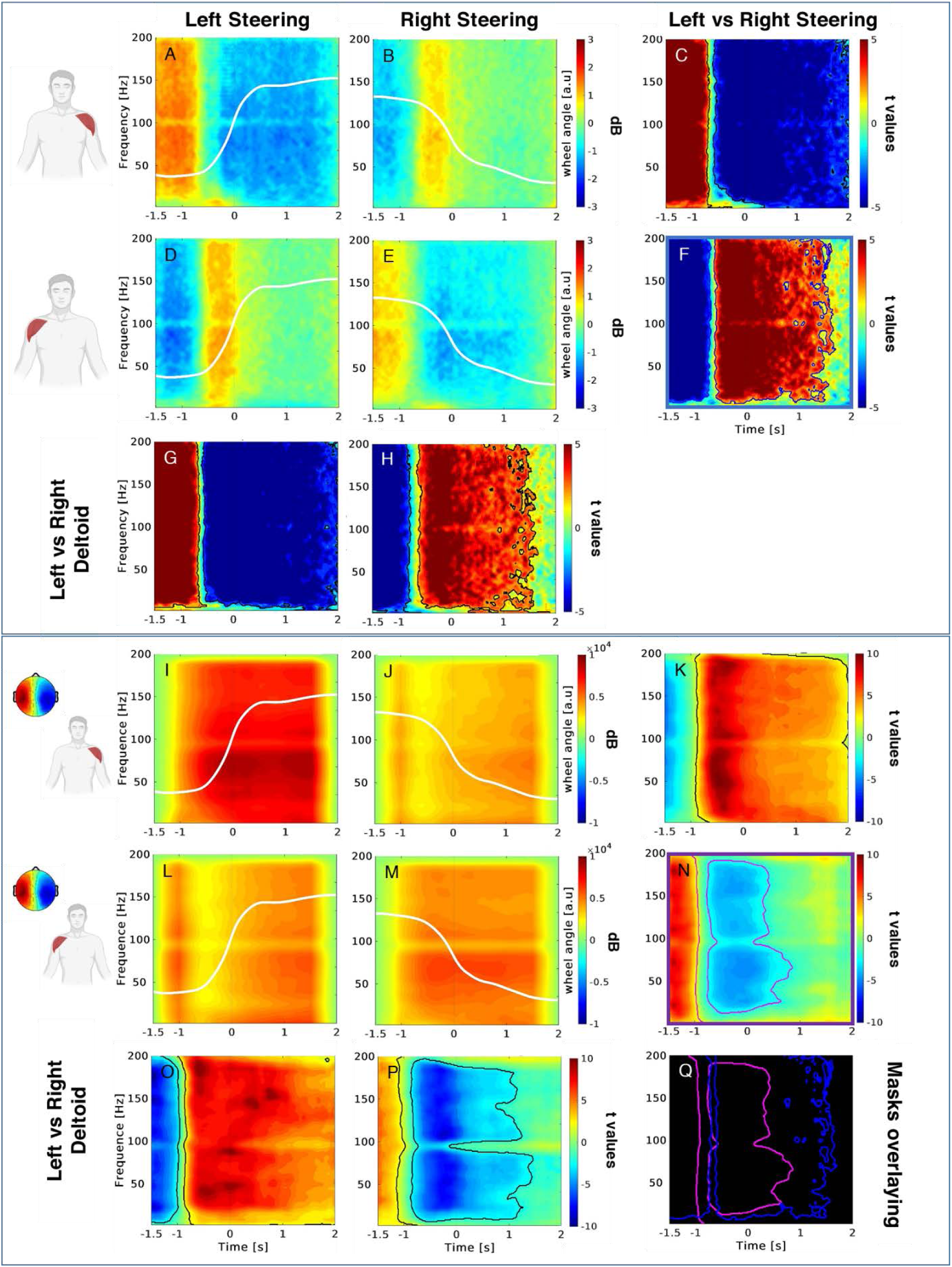
ERSP for the EMG signals collected from the deltoids during the non-ecological steering task (upper box), and cross-correlation results between EEG and EMG data (lower box). Upper box. First and second rows (from the top) illustrates the EMG ERSP for the left and right deltoid during left (A,D) and right (B,E) steering as well as the statistical comparison of the two conditions (C,F). Third row illustrates the statistical comparisons of the EMG ERSP between left and right deltoid during left (G) and right (H) steering. Colorbars indicate in red (blue) the synchronization (relaxation) of the EMG activity (A, B, D, E) with respect to the baseline, as well as the increase (decrease) of such activity during the left (right) steering for the right (left) deltoid, as well as for the left (right) deltoid during right (left) steering (C, F, G, H). Lower box. First and second rows (from the top) illustrates the EEG-EMG cross-correlation values for the left and right deltoid during left (I,L) and right (J,M) steering as well as the statistical comparison of the two conditions (K,N). Third row illustrated the statistical comparisons of the EEG-EMG cross-correlation values between left and right deltoid during left (O) and right (P) steering. Colorbars indicate in red (blue) the increase (decrease) of the EEG-EMG cross-correlation values (I, J, L, M), as well as the increase (decrease) of such values during the left (right) steering for the right (left) deltoid, as well as for the left (right) deltoid during right (left) steering (K, N, O, P). White lines depict the left and right steering wheel angle profiles (A, B, D, E, I, J, L, M). Contour mask (C, F, G, H, K, N, O, P) delimits the statistically significant portion of the corresponding compared panels (non-parametric t-test, cluster corrected). Panel Q highlights the significant masks corresponding to the comparisons left versus right steering for the right deltoid for the EMG ERSP data (blue contours, F) and EEG-EMG cross-correlation (magenta contours, N)

The lower box of Figure 3 shows the results related to the cross-correlation analysis performed between the EEG mask gathered during the non-ecological task and the EMG ERSP panel corresponding to the ecological scenario in the same steering condition and for each of the two deltoids (i.e., non-eco EEG in left steering cross-correlated with the eco EMG in left steering). Here we can observe a higher cross-correlation values associated to the left deltoid during left steering (panel I) and for the right deltoid during the right steering (panel M) when compared with the right (panel J) and left (panel L) steering, respectively. This is statistically demonstrated with the corresponding non-parametric analysis (panel K, N). Moreover, such differences between cross-correlation values is also observed during left (panel O) and right (panel P) steering when comparing the left and right deltoid. Strikingly, these statistics show that the masks of significant activations related to the cross-correlation analysis are detected earlier (magenta contour in panel Q) with respect to the activations only EMG ERSP (blue contour in panel Q), for the condition right deltoid, left versus right steering.

Figure 4 shows the results related to the cross-correlation analysis performed between the same EEG mask gathered during the non-ecological task and the EMG ERSP panels corresponding to the ecological scenario in the non-steering (upper box) and shuffled steering (lower box) condition. Here we can observe low cross-correlation values associated to all conditions which were reported on the same scale of Figure 3 (lower box). Finally, the corresponding statistics returned no significant difference for any condition.

**Fig.4.**
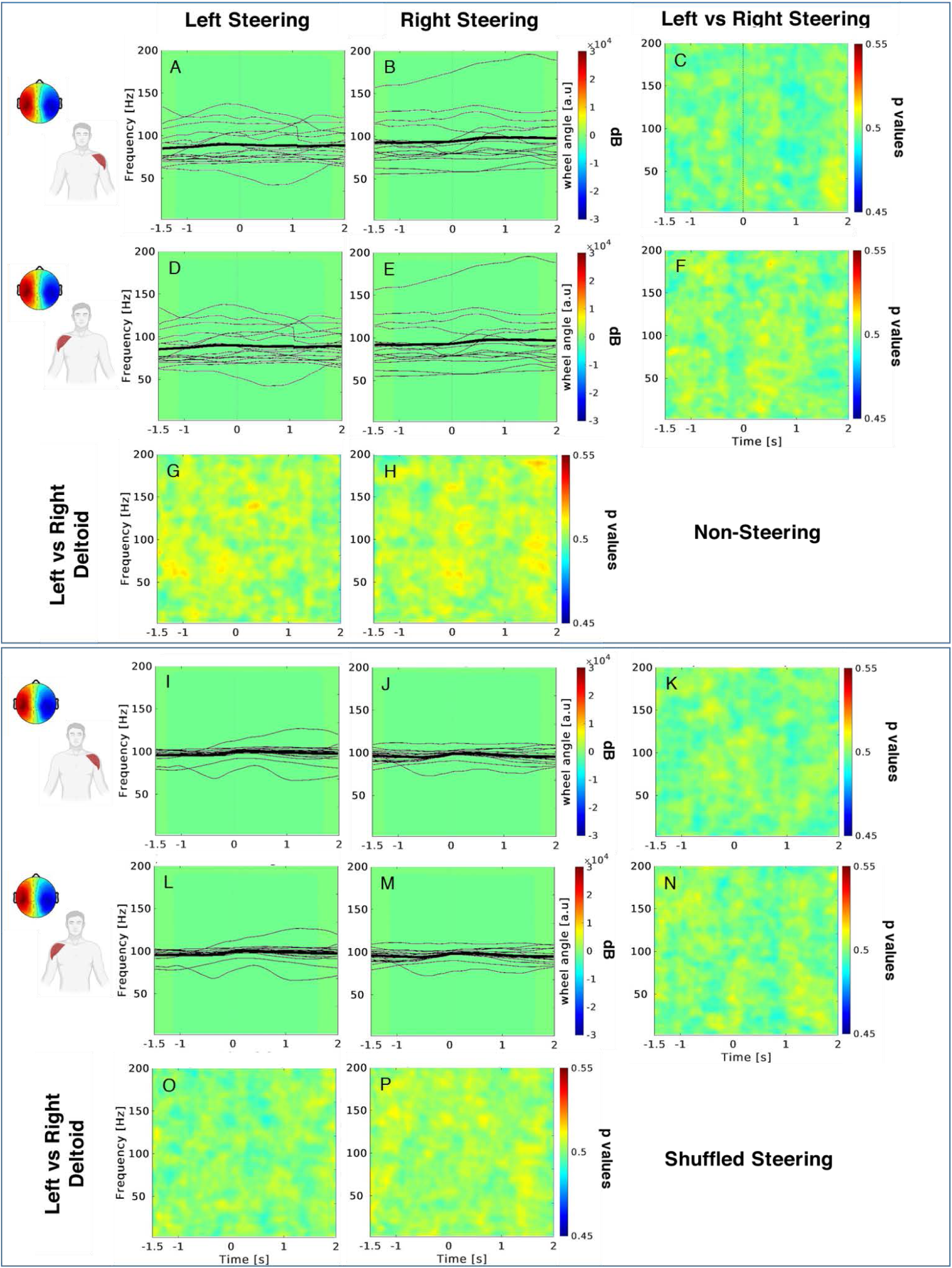
Cross-correlation results between EEG and EMG data related to non-steering (upper box) and shuffled steering (lower box) conditions. Same convention as Figure 3 (lower box). Black lines depict the left and right steering wheel angle profiles as butterfly plot. Thick lines represent the average across subjects (thin lines)

Variations of the wheel angle for the left and right steering were represented with the white (Figure 3) and black (Figure 4) signals within each corresponding panel.

## 4. Discussion

In the present study we report that the modulation of the EEG mu rhythm observed during the motor preparation of non-ecological steering predicts the muscular activity of deltoids, thus anticipating subject steering behaviour. The reactivity of such rhythm measured across sensorimotor areas during the non-ecological steering preparation anticipates the corresponding action. We report the increase of EMG activity of the deltoid anticipating the contralateral steering in non-ecological and ecological steering tasks. These results show an asymmetric muscular activity of the deltoids beginning before the action onset remaining steady during the steering execution, i.e. the coordinated increase of power of the right deltoid and the corresponding decrease of power of the left deltoid are associated to the steering action on the left side. Such findings show that by monitoring the muscular activity of the two deltoids it is possible to discriminate the steering side before the action onset while driving in non-ecological and ecological scenarios. Strikingly, the identified non-ecological EEG feature correlates with the ecological EMG activity of the deltoids, providing an improvement of the discrimination power of the steering side during driving simulation. The comparison between the results related to the EMG analysis and those concerning the EEG-EMG cross-correlation returned larger relative timing in favour of the latter related to the succession of steering events. In fact, from one side there is the issue to distinguish the anticipatory components due to EEG predicting power from those which are merely due to computational accounts, e.g. windowing implied in the cross-correlation calculation. However, we do observe a clear anticipatory pattern returned by cross-correlation values which is related to the succession between left and right steering events. It is just this change of cross-correlation values which is detectable in advance with respect to the single EMG signal.

The timing of the EEG response identified in this study is compatible with patterns of event related desynchronization (ERD) which are reported to be predictive of the upcoming action between 1.5 and 1 second prior the movement onset (Neuper et al. 2006). ERD reflects changes of non-phase-locked EEG oscillatory activity within the alpha and beta frequency due to sensorimotor events (Pfurtscheller and Lopes da Silva 1999; Savić et al. 2020). The EEG component identified during the non-ecological task arises from the motor areas and may reflect the preparation of steering actions. The scalp topography that we reported is reminiscent of the mu rhythm observed in motor and premotor regions (Gastaut et al. 1952; Pfurtscheller et al. 1997). Such electrical activity is known to produce somatotopically organized desynchronization during execution, observation and imagination of actions (Pineda 2005; Pfurtscheller et al. 2006; Arnstein et al. 2011; Avanzini et al. 2012). These regions perform several functions other than control of body movements, such as sensory-motor transformation, action understanding, decision processing regarding execution and initiation of action, preparation and planning of complex movements (Roland 1984; Rizzolatti and Luppino 2001). Recent literature reports ERD in contralateral sensorimotor cortices during movement preparation of visually cued movements (Li et al. 2018; Little et al. 2019). Therefore, we argue that the identified EEG mu rhythm modulations regulate the motor preparation of the right and left deltoids for steering actions. In particular, we report that the mu desynchronization over left sensorimotor regions is related with the activity of the right deltoid during left steering. This would show that left steering involve larger neural computation with respect to right steering, despite the absence of significant difference in the steering wheel angle, as already reported in a previous study (Oka et al. 2015). Another study also reports the desynchronization of the alpha rhythm across sensorimotor regions related to relative steering angle compensation (Brooks and Kerick 2015), thus showing the relation between such electroencephalographic feature and steering response as already observed in more simple tasks (Pfurtscheller and Neuper 1994; Stancák and Pfurtscheller 1996).

By means of the analysis of the EMG in the time-frequency domain we observe that the maximum muscle activity changes significantly due to different steering wheel angle and turning direction in both non-ecological and ecological scenarios. It is known that the anterior and middle portions of the deltoid muscle work intermittently during car driving and that their functioning regulate the contralateral rotation of the steering wheel, being activated for a duration of around 50% since the initial stage of the action (Jonsson and Jonsson 1975; Pick and Cole 2006; Liu et al. 2012; Gao et al. 2014). Specifically, when the steering wheel is near its centre, muscle activity is relatively small but it increases rapidly as the wheel starts to turn (Gao et al. 2014). Although EMG was successfully used to identify the muscles involved in generating and predicting torque at the steering wheel (Pick and Cole 2006), EMG alone has a low power in predicting the steering as we report to be limited to a few hundreds of milliseconds. Thus, we exploited the EEG scalp feature to improve the detection of steering by computing the cross-correlation between the mu rhythm desynchronization retrieved during the non-ecological task and the EMG activity elicited in the ecological scenario. This procedure allowed us to enlarge the time window in which it is possible to discriminate the steering action. We showed that this EEG-EMG coupling is specific both for steering action and for its directionality and returned better results when comparing with single EMG signals as also recent literature reports in hybrid human-machine interface for gait decoding (Tortora et al. 2020), motor rehabilitation of stroke patients (Sarasola-Sanz et al. 2017), movement detection of hand-paralysed patients (Lóopez-Larraz et al. 2018) and further fields for the general aim to improve classification accuracy and to increase the number of commands for a variety of Human-Machine-Interfaces (Hong and Khan 2017).

Here we performed two consecutive recording sessions to demonstrate the feasibility to exploit EEG correlates of steering behaviour collected in a standard highly controlled environment to detect steering action in a more ecological setting such as the one of a driving simulator. In the first non-ecological recording session we extracted ERD as a neural correlate of the EMG activity of the deltoids predicting the upcoming steering behaviour. In the following ecological recording session performed at the driving simulator we collected the EMG activity of the deltoids and showed that the ERD resulted from the previous session is indeed informative concerning the steering action in this more naturalistic scenario.

Several recent studies on driving addressed the issue of steering classification by means of different EEG features. In particular, it was assessed the possibility to decode self-generated actions detecting whether the driver would perform a lane change in the close future in a simulated highway (Gheorghe et al. 2013). Authors report slow negative EEG deflections across central areas which are consistent with the movement-related potentials 500 ms before the lane change, yielding classification accuracy of 79% with an average detection time of 613 ms before the actual steering action. In another series of study performed with both driving simulator and real car (Zhang et al. 2015), error-related brain potentials were analysed to investigate the possibility to use an external device to be adapted to the driver’s goal, i.e. assisting in the upcoming steering action. This was realized by showing the drivers a visual stimulus that indicates its inference about the next direction of turning when approaching an intersection. Authors report differences in the EEG response over fronto-central areas when the directional stimulus does not match the driver’s intention. Statistical differences between error and correct conditions were observed between 200 ms and 600 ms after feedback, yielding a mean accuracy of the event-related decoding of 0.68 which indicates the possibility to extract meaningful information about the driver’s need of assistance.

The existence of preparatory electrophysiological activity elicited before the onset of steering action allows to infer the upcoming driving actions in advance. The reported EEG-EMG coupling is a proof of concept for utilizing hybrid systems for the detection and online prediction of driving actions, exemplifying how it might be possible to complement information from behavioral, physiological, and external sources to control the level of assistance needed by the driver in that context (Chavarriaga et al. 2018). The predictive power of the EEG-EMG coupling demonstrated in a car simulator could be further investigated in a larger sets of actions to extend the validity of this neurophysiological mechanism beyond driving.

## Declarations

### Availability of data and material

The data that support the findings of this study are available from CAMLIN Limited but restrictions apply to the availability of these data, which were used under licence for the current study, and so are not publicly available. Data are however available from the authors upon reasonable request and with permission of CAMLIN Limited.

### Code availability

Custom code is available upon reasonable request.

### Authors’ contributions

All authors contributed to the study conception and design. Material preparation and data collection were performed by Giovanni Vecchiato and Maria Del Vecchio. Data analysis was performed by Giovanni Vecchiato. The first draft of the manuscript was written by Giovanni Vecchiato and all authors commented on previous versions of the manuscript. All authors contributed and approved the final manuscript.

### Ethics approval

The research was approved by the local Ethical Committee (comitato etico Unico per la Provincia di Parma).

### Consent to participate

All subjects expressed written consent to participate in the research.

### Consent for publication

All Authors agreed on the final version of the manuscript.

## Acknowledgements

The present study was supported by a research agreement between CAMLIN Limited and IN-CNR, and a research agreement between CAMLIN Limited and Toyota Motor Europe.

